# PFKFB3 inhibition restores β-cell responsiveness to acetylcholine in a 3xTg-AD pancreatic tissue-slice model

**DOI:** 10.64898/2026.09.30.755766

**Authors:** Johannes Pfabe, Sandra Postic, Lea Miglinci, Anita Krnjic, Hifza Ahmed, Tobias Eisenberg, Jelena Tadic, Kavit Raval, Gurprit Bhardwaj, Slavica Tudzarova, Marjan Slak Rupnik

## Abstract

The glycolytic regulator PFKFB3 supports dysfunctional β-cell states, but its role in pancreatic Ca^2+^ signaling remains unclear. We examined responses to PFKFB3 inhibitors (AZ67, AZ26) in pancreatic tissue slices from aged 3xTg-AD mice, a model of Alzheimer’s disease that develops impaired glucose tolerance and diabetes as comorbidity. Compared to wild-type controls, β-cells showed weaker glucose-stimulated and acetylcholine-potentiated Ca^2+^ activity, accompanied by increased nuclear PFKFB3 positivity in insulin-defined islet-cell populations, used here as a PFKFB3-associated stress-state marker. During glucose and acetylcholine stimulation, AZ67 increased β-cell Ca^2+^ event frequency while maintaining short events. In neighboring acinar cells, PFKFB3 inhibitor exposure was followed by renewed oscillations after acetylcholine-evoked activity had waned. β-cell recruitment was preserved during adrenergic inhibition, and enhanced activity persisted after inhibitor withdrawal, although these recordings were exploratory and residual tissue drug exposure could not be excluded. An effect that survives both drug removal and adrenergic opposition identifies a Ca^2+^ signaling phenotype in 3xTg-AD β-cells that is acutely inducible yet not transient, and establishes PFKFB3 as a regulator of muscarinic responsiveness in the stressed β-cell.

**graphical abstract:** 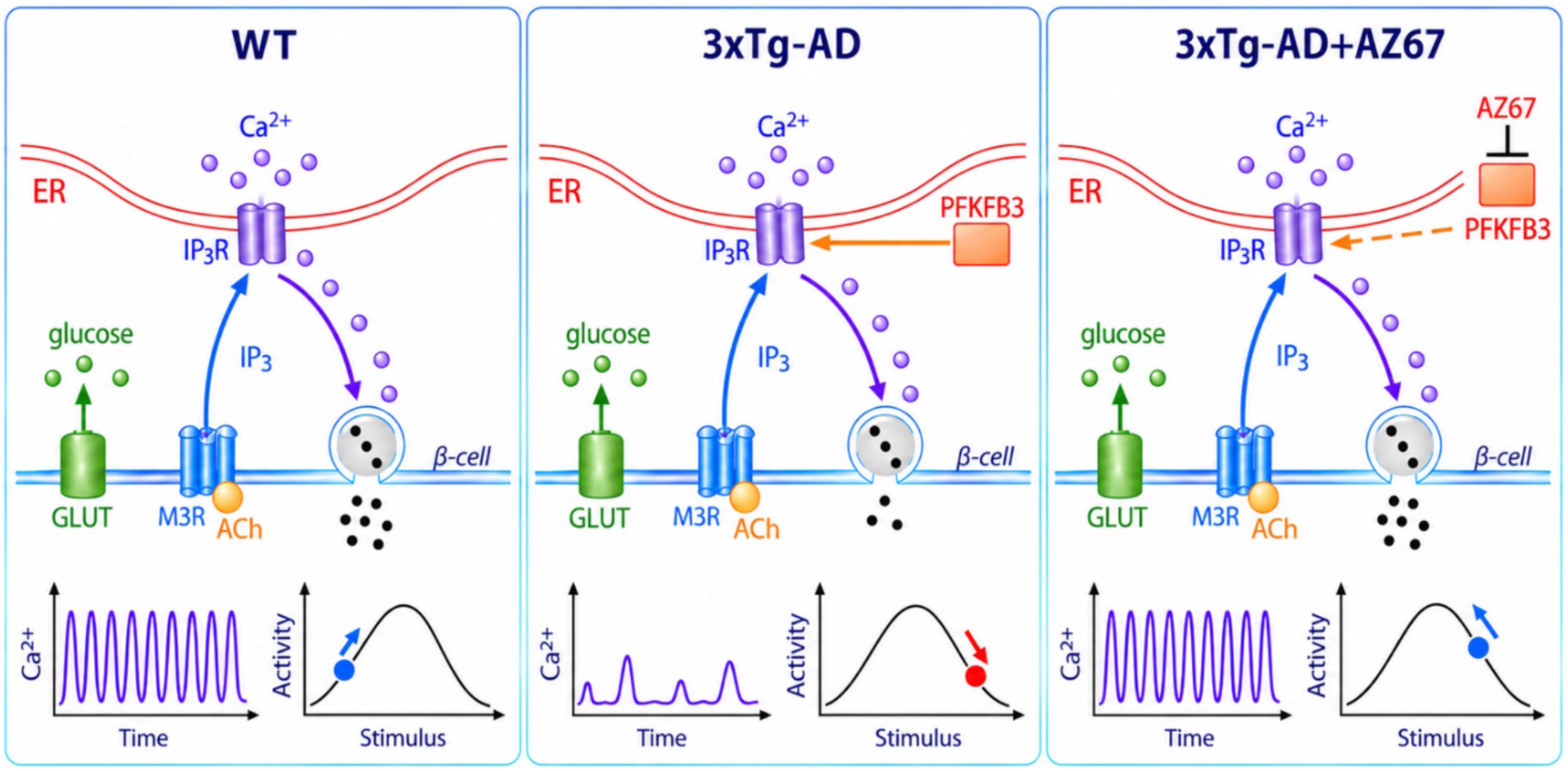

**Graphical abstract: PFKFB3 inhibition enhances muscarinic β-cell Ca^2+^ responsiveness in 3xTg-AD pancreatic tissue slices.** β-cells in 3xTg-AD pancreatic slices exhibit lower glucose-stimulated and acetylcholine (ACh)-potentiated Ca^2+^ activity than wild-type (WT) controls. During combined glucose and ACh stimulation, AZ67 increases β-cell Ca^2+^ event frequency while maintaining a short-event phenotype. The bell-shaped curves illustrate a working model in which PFKFB3-associated dysregulation shifts muscarinic/IP_3-_linked signaling beyond an optimal range for oscillatory activity, whereas PFKFB3 inhibition favors a return toward this range. The schematic thus distinguishes enhanced Ca^2+^ oscillatory activity from a simple increase in Ca^2+^ release. Direct regulation of β-cell IP_3 r_eceptors by PFKFB3 and the depicted consequences for insulin secretion remain proposed mechanisms and functional implications, respectively, rather than outcomes established in these slice experiments.

## INTRODUCTION

Inositol 1,4,5-trisphosphate receptors (IP₃Rs) mediate Ca^2+^ release from the endoplasmic reticulum (ER) downstream of Gq-coupled receptor activation and contribute to stimulus-secretion coupling in pancreatic cells^1,2^. In acinar cells, acetylcholine (ACh) and cholecystokinin evoke polarized Ca^2+^ oscillations that drive digestive enzyme secretion^2–4^. In pancreatic β-cells, muscarinic signaling potentiates glucose-stimulated activity and insulin secretion through phospholipase C activation, IP₃ production, and IP₃R-dependent Ca^2+^ release that complements voltage-gated Ca^2+^ entry^5–7^. Although both populations share IP₃R signaling machinery, differences in receptor isoforms, subcellular localization, and downstream functions shape their distinct Ca^2+^ responses^8,9^.

Metabolic state can directly influence Ca^2+^ signaling^10^. Several glycolytic enzymes associate with IP_3R_s and modulate receptor gating, linking metabolic activity to intracellular Ca^2+^ release^11,12^. The glycolytic regulator 6-phosphofructo-2-kinase/fructose-2,6-bisphosphatase 3 (PFKFB3) has been recently shown to regulate IP₃R function in acinar cells and promote pathological Ca^2+^ overload during pancreatitis^13^. In β-cells, PFKFB3 is induced under proteotoxic, metabolic, and inflammatory stress^14–16^. This response can support stressed-cell survival at the expense of differentiated function, allowing dysfunctional β-cell states to persist^14,17^. β-cell specific disruption of PFKFB3 has been reported to restore cell-fitness competition under diabetogenic stress^16^, while pharmacological inhibition improves glucose tolerance and glucose-stimulated insulin secretion, with benefits persisting after treatment withdrawal^18^. These findings identify PFKFB3 as a potential link between stress adaptation and impaired β-cell responsiveness.

The functional consequences of PFKFB3 inhibition may nevertheless differ between pancreatic compartments. In acinar cells, PFKFB3 promotes excessive IP₃R-mediated Ca^2+^ release, whereas its induction in stressed β-cells accompanies impaired stimulus–secretion coupling^14,15^. Its inhibition could therefore reshape Ca^2+^ signaling in a cell-type- and state-dependent manner. Whether it acutely improves muscarinic responsiveness in dysfunctional β-cells remains unresolved. We addressed this question using PFKFB3 inhibitors AZ67 and AZ26^19^ during sequential glucose–ACh stimulation, with simultaneous assessment of β-cell and neighboring acinar cell activity. We also examined nanomolar AZ67 exposure, because nominal concentrations required in intact-tissue may differ from biochemical potency owing to compound binding, partitioning and diffusion^20^. To investigate these responses, we used pancreatic tissue slices from aged 3xTg-AD mice with impaired β-cell glucose responsiveness^21^. Although developed as a model of Alzheimer’s disease^22^, multiple studies reported age-dependent disturbances in glucose homeostasis and pancreatic amyloidopathy^23–25^, making this pancreatic phenotype relevant to impaired glucose tolerance and diabetes as metabolic comorbidities of AD^26^. This setting provides a complementary context to human islet amyloid polypeptide-transgenic mice maintained on high-fat diet (hTG+HFD), in which amyloid-associated proteotoxic stress engages the PFKFB3 pathway^14–16^. Human IAPP and amyloid-β share structural features within amyloid-forming regions and can interact and cross-seed under experimental conditions^27,28^. These similarities suggest that distinct amyloidogenic proteins may engage convergent stress responses in pancreatic cells. Our focus is whether the β-cell dysfunction present in 3xTg-AD model retains an acutely modifiable Ca^2+^ signaling component.

In addition to metabolic and cholinergic stimulation, β-cell activity is constrained by sympathetic input. Epinephrine inhibits β-cell electrical activity, Ca²⁺ signaling and insulin secretion, predominantly through α₂-adrenergic signaling^29^. We used this physiological inhibitory input to benchmark AZ67 responses in WT tissue and to examine β-cell recruitment during an opposing adrenergic challenge in 3xTg-AD slices. Because a companion study reports benefits persisting after treatment withdrawal, we also explored whether altered Ca²⁺ activity remained detectable after removing AZ67 or AZ26 from the perfusate.

Here, we combined pancreatic-slice Ca²⁺ imaging with PFKFB3 immunostaining to characterize impaired β-cell responsiveness in aged 3xTg-AD mice and its acute pharmacological modulation. Neighbouring acinar cells provided a simultaneous comparison of endocrine and exocrine responses, while adrenergic challenges and prolonged recordings explored the maintenance of β-cell activity.

## RESULTS

### β-cells in 3xTg-AD mice show weakened glucose responsiveness and lower Ca^2+^ activity during ACh stimulation

To assess glucose-dependent β-cell Ca^2+^ signaling in situ, we recorded cytosolic Ca^2+^ dynamics in acute pancreatic tissue slices in response to glucose stimulation. We employed a stepwise glucose ramp protocol spanning the physiological activation range using established pancreatic-slice Ca^2+^ imaging and event-detection workflows^30,31^. In slices obtained from both genotypes (WT and 3xTg-AD), β-cells initiated oscillatory activity after crossing a threshold near ∼6.5 mM glucose (Fig. 1A-C). Glucose significantly increased Ca^2+^ event AUC in WT islets (positive main effect of glucose; β = 0.914, p < 0.001). However, a significant genotype x glucose interaction (β = −0.312, p < 0.01) indicated that β-cells from 3xTg-AD pancreatic slices exhibited a weaker glucose dependence compared to WT controls (Fig. 1C), consistent with impaired metabolic coupling. Slice-level analyses retained the substantial biological heterogeneity among preparations, consistent with our previous findings in a larger dataset, while an animal-level sensitivity analysis for Fig. 1C also supported reduced glucose dependence in 3xTg-AD β-cells (β = −0.303, p < 0.05). For the glucose-ramp analysis (Fig. 1A–C): WT, n = 6 mice, 6 slices; 3xTg-AD, n = 4 mice, 5 slices. For the ACh analysis (Fig. 1D–F): WT, n = 6 mice, 6 slices; 3xTg-AD, n = 4 mice, 17 slices.

**Figure 1.**
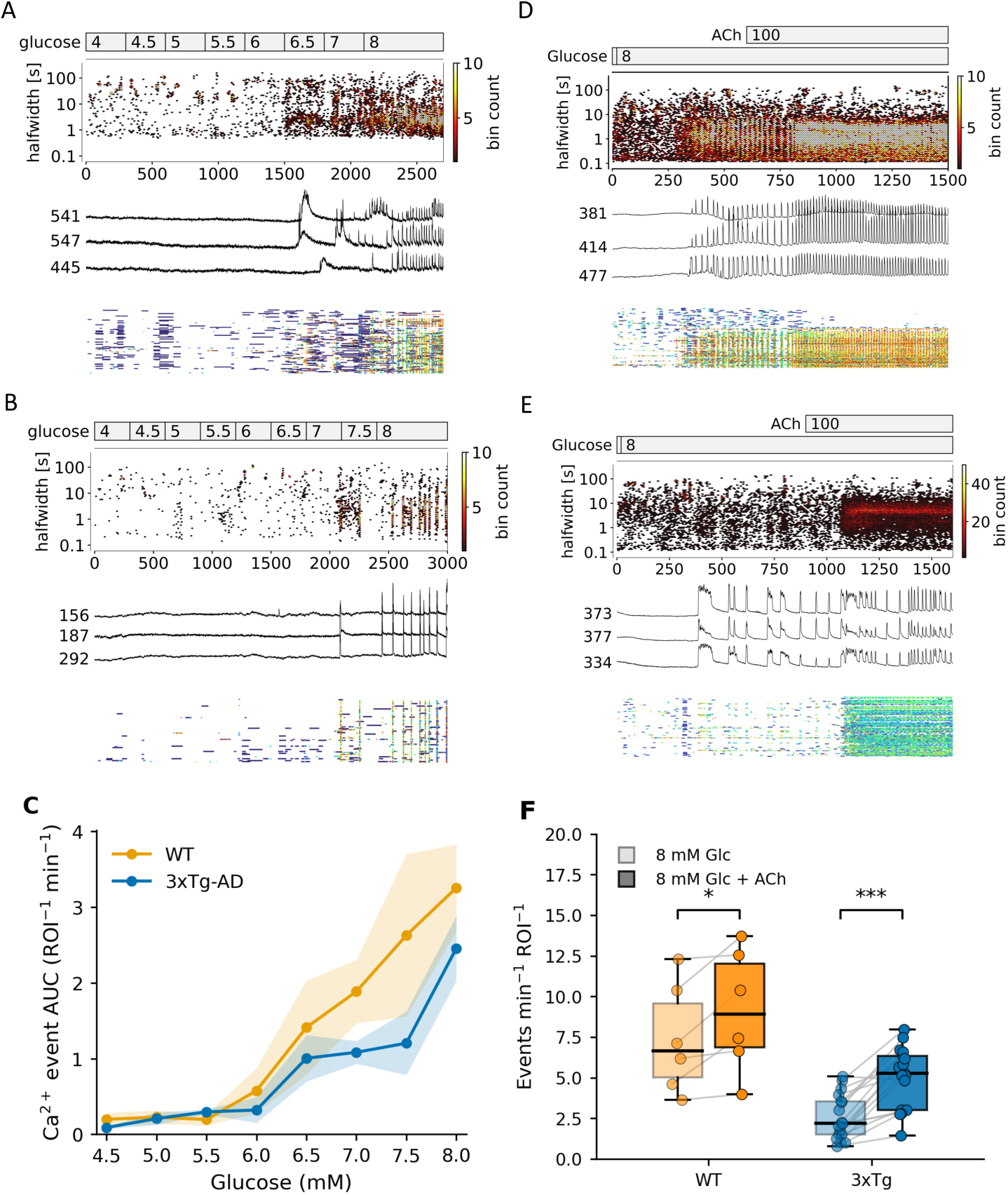
Reduced glucose-dependent and Ach-stimulated Ca^2+^ activity in 3xTg-AD β-cells. (A, B) Representative recordings from WT (A) and 3xTg-AD (B) pancreatic tissue slices during a stepwise glucose ramp from 4 to 8 mM). Each panel shows a time-resolved distribution of Ca^2+^ event halfwidths, representative single-ROI traces and a multi-ROI event raster. Time and event halfwidth are expressed in seconds. Color scales indicate event counts per bin, and numbers beside traces identify individual ROIs. (C) Ca^2+^ event AUC per ROI per minute across glucose concentrations in WT (orange) and 3xTg-AD (blue) islets. Lines and shading denote mean ± SEM. A linear mixed-effects model identified a reduced glucose dependence in 3xTg-AD β-cells (genotype × glucose interaction, β = −0.312, p < 0.01). WT: n = 6 mice, 6 slices; 3xTg-AD: n = 4 mice, 5 slices). (D, E) Representative responses to addition 100 nM ACh at constant 8 mM glucose, in WT (D) and 3xTg-AD (E) islets. (F) β-cell event frequency, expressed as events min^−1^ ROI^−1^, before and after ACh addition. Box plots show medians and interquartile ranges. Points represent individual islets, with paired observations connected. Brackets indicate within-genotype comparisons, between genotype comparisons are reported in the Results. WT: 6 mice, 6 slices; 3xTg-AD: 4 mice, 17 slices. *p < 0.05; ***p < 0.001.

We next tested muscarinic potentiation by adding ACh (100 nM) during glucose-stimulated activity. ACh increased β-cell Ca^2+^ oscillation frequency in both genotypes (paired comparisons: WT, p = 0.036; 3xTg-AD, p < 0.001), whereas frequency was lower in 3xTg-AD than in WT both at 8 mM glucose alone (p=0.032) and after addition of 100 nM ACh, p=0.044, Fig. 1D-F). In the same slices, ACh activated the majority of acinar cells, and acinar activation was broadly comparable between genotypes, indicating that the major genotype-dependent deficit was observed in β-cells (see below).

### ACh evokes bell-shaped Ca^2+^ responses in β-cells and acinar cells

As previously reported, muscarinic stimulation can exert both stimulatory and inhibitory effects on pancreatic Ca^2+^ signaling depending on agonist concentration^6,32^. We therefore quantified the concentration dependence of ACh responses in pancreatic slices C57B6/J WT mice (more than 10 months old) (Fig. 2). In β-cells, ACh concentration significantly affected Ca²⁺ event frequency (repeated-measures ANOVA, p < 0.001), with event frequency increasing from 0 to 100 nM ACh (Holm-adjusted paired comparison, p < 0.01) and declining again at higher concentrations from 250 to 500 nM ACh (Holm-adjusted paired comparison, p < 0.01) (Fig. 2B). Event halfwidth did not differ significantly across ACh concentrations (RM-ANOVA, p = 0.518, Fig. 2C). A one-sided inverted-U contrast analysis provided strong evidence for a bell-shaped dependence (p < 0.001).

**Figure 2.**
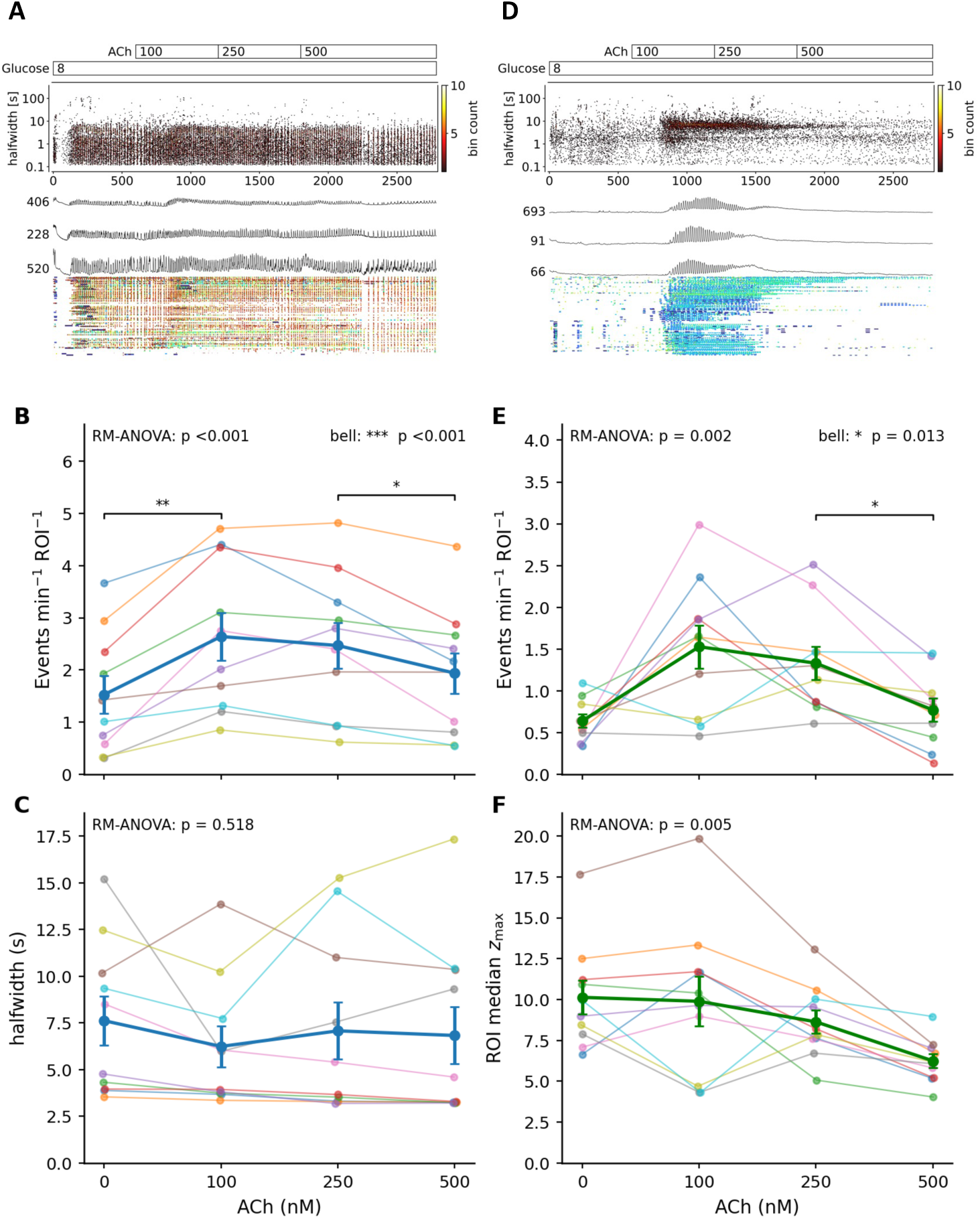
Bell-shaped dependence of pancreatic Ca^2+^ event frequency on ACh concentration. (A,D) Representative β-cell (A) and acinar cell (D) recordings during sequential exposure to 0, 100, 250, and 500 nM ACh at 8 mM glucose. Panels show time-resolved event-halfwidth distributions, representative single-ROI traces and multi-ROI event rasters. Time and halfwidth are expressed in seconds. Color scales indicate event counts per bin. (B,C) β-cell event frequency (B, events min^−1^ ROI^−1^) and halfwidth (C, seconds). (E,F) acinar cell event frequency (E, events min^−1^ ROI^−1^) and event amplitude (F, ROI median event z_max, dimensionless). Thin lines connect measurements from individual slices. Bold lines, symbols and error bars indicate the group mean ± SEM. ACh concentration effects were assessed by repeated-measures ANOVA (RM ANOVA), followed up by paired two-sided t-tests with Holm correction. A one-sided inverted-U contrast tested whether the event frequency at 100 nM ACh exceeded the mean frequency at 0 and 500 nM ACh. Its significance is indicated as “bell” in B and E. 3 male mice, 10 slices. *p < 0.05, **p < 0.01 and ***p < 0.001.

Acinar cells showed a closely related concentration dependence. Event frequency increased strongly at 100 nM ACh and subsequently declined at 250 and 500 nM (repeated-measures ANOVA, p < 0.002), with significant evidence for a bell-shaped frequency response (p < 0.013; Fig. 2E). Event amplitude, quantified as ROI median z-score, also varied with ACh concentration (repeated-measures ANOVA, p = 0.028), with the lowest values observed at 500 nM ACh (Fig. 2F). Thus, despite their distinct Ca^2+^ signaling phenotypes, both β-cells and acinar cells display a concentration-dependent transition from stimulation at lower ACh concentrations toward inhibition at higher concentrations. The non-monotonic ACh response provides a rationale for considering whether reduced muscarinic/IP_3-_linked drive could restore oscillations under some conditions. It does not by itself establish the position of 3xTg-AD β-cells on an IP_3R_ response curve or identify the mechanism of AZ67 action.

### 3xTg-AD islets show nuclear-selective PFKFB3 upregulation across insulin-defined cell strata

To relate the functional Ca^2+^ phenotype to a stress associated β-cell state, we quantified PFKFB3 immunoreactivity at single-cell resolution in pancreatic sections (Fig. 3). Using automated threshold (mean, mean+1 SD and mean+2 SD) -based thresholds to classify nuclear and cytoplasmic PFKFB3 intensity, and stratifying islet cells by insulin expression (INS^high^, INS^low^,INS^−^), we compared WT and 3xTg-AD pancreata and the PFKFB3 immunostaining against tissue from human IAPP-transgenic mice maintained on a high-fat diet (hTG+HFD), an established model of β-cell proteotoxicity and metabolic stress and a positive control for PFKFB3⁺ β-cell dysfunction^14–16^. 3xTg-AD pancreata also showed a shift in overall islet cell composition, with a reduced INS^high^ fraction and an increased INS^−^ fraction relative to WT (Fig. 3A-D)^33,34^. 3xTg-AD islets showed a significant increase in nuclear PFKFB3 positivity relative to WT across all three insulin-content strata examined (Fig. 3G-I). In contrast, cytoplasmic PFKFB3 did not differ significantly between genotypes (Fig. 3F). Together, these findings identify increased nuclear PFKFB3 positivity across insulin-defined islet-cell populations in aged 3xTg-AD mice. We interpret this nuclear signal as a PFKFB3-associated stress-state marker providing tissue-level context for the pharmacological experiments that follow, rather than as evidence of the effector pool responsible for the acute Ca²⁺ phenotype.

**Figure 3.**
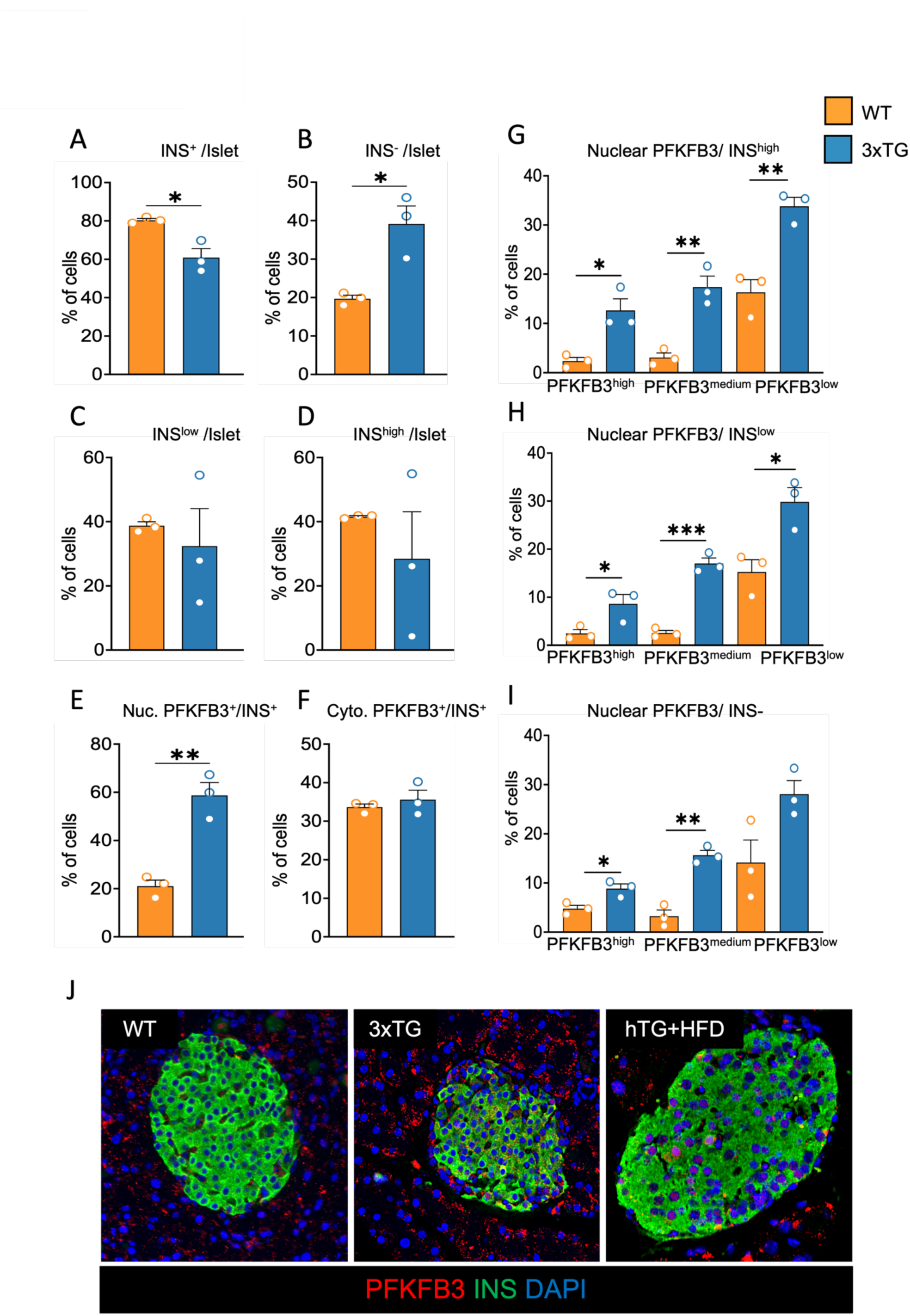
Compartment-specific PFKFB3 upregulation in 3xTg-AD islets. (A–D) Quantification of insulin-positive (INS⁺, A), insulin-negative (INS^−^, B), insulin-low (INS^low^, C), and insulin-high (INS^high^, D) cell fractions as a percentage of total islet cells in WT and 3xTg-AD pancreata. (E, F) Quantification of PFKFB3 immunoreactivity in INS⁺ cells, comparing nuclear (E) and cytoplasmic (F) PFKFB3-positive fractions between WT and 3xTg-AD. Cells were binned into PFKFB3 high/medium/low/negative categories using automated thresholds derived from the single-cell intensity distribution (see Methods). (G–I) Nuclear PFKFB3 classification (high/medium/low) within each insulin-defined stratum: INS^high^ (G), INS^low^ (H), and INS^−^ (I) in WT and 3xTg-AD islets. (J) Representative immunofluorescence images of pancreatic sections from WT and 3xTg-AD mice immunostained for PFKFB3 (red), insulin (green), and nuclei (DAPI, blue). Pancreata from hTG+HFD-transgenic mice were processed and stained in parallel as a positive control. 3xTg-AD islets show an increase in nuclear PFKFB3-positive cells relative to WT, consistent across all three insulin-defined strata (G–I), while cytoplasmic PFKFB3 (F) did not differ significantly between genotypes. Across all three insulin strata, 3xTg-AD islets also show a shift in overall islet composition, with a reduced INS^high^ fraction and increased INS^−^ fraction relative to WT (A–D). Data are presented as mean ± SEM (n = 3 mice per group; 20–35 islets analyzed per pancreas). Statistical analysis was performed using an unpaired t-test. Statistical significance is indicated as follows: P < 0.05 (*), P < 0.01 (**), and P < 0.001 (***)

### Age-matched WT control experiments distinguish AZ67 from physiological adrenergic inhibition

To confirm the stability of the preparation for prolonged pharmacological protocols, we first assessed the effect of the vehicle (2-hydroxypropyl-β-cyclodextrin, 2-HPBCD) on ACh-stimulated slices from WT mice. Vehicle exposure did not significantly alter the frequency or duration of β-cell Ca^2+^ oscillations and did not affect the characteristic time-dependent rundown of ACh-evoked activity in acinar cells (Fig. 4A, D).

**Figure 4.**
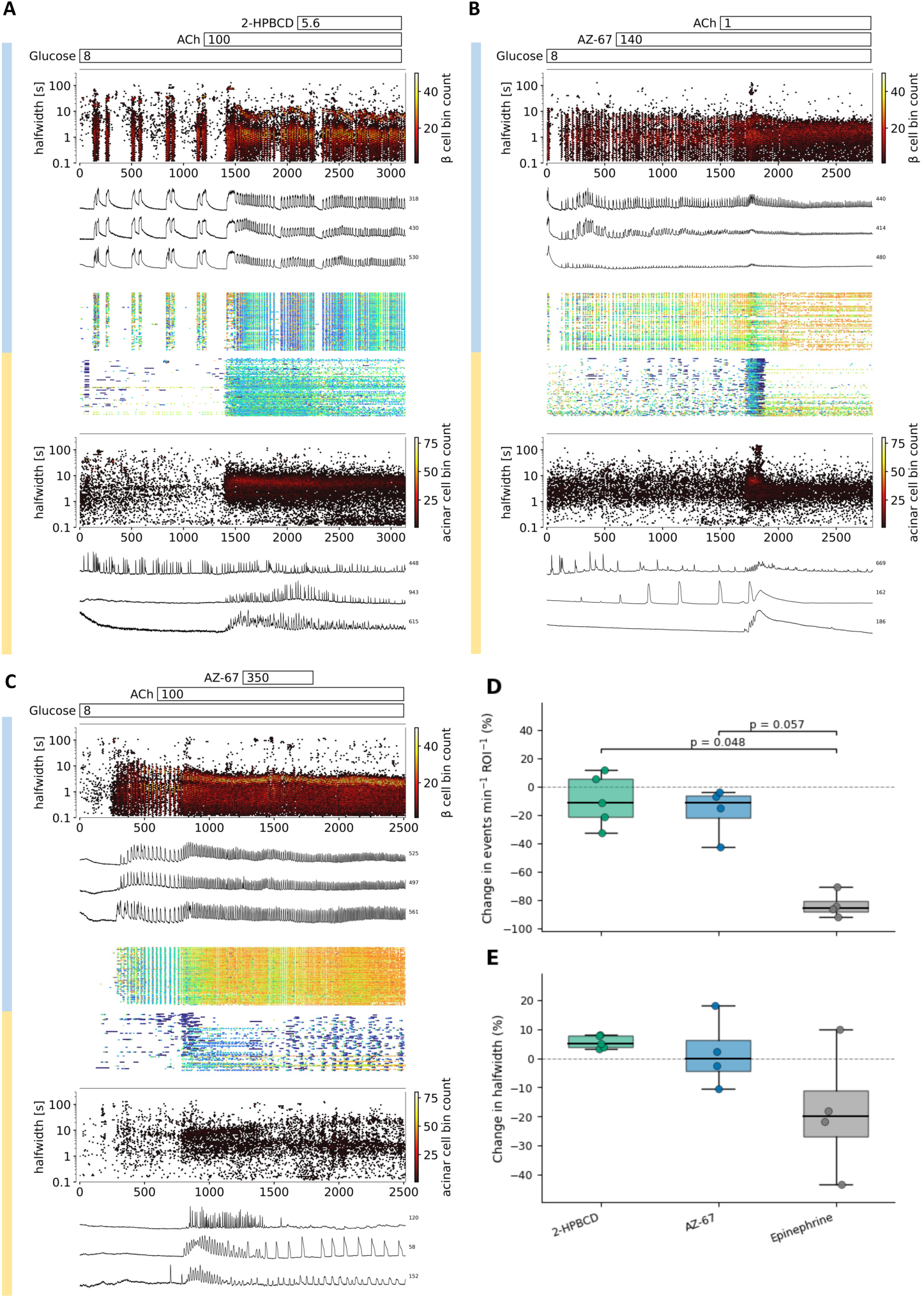
Vehicle, AZ67 and epinephrine responses in WT pancreatic slices. (A-C) Representative recordings showing vehicle exposure (2-hydroxypropyl-β-cyclodextrin, 2-HPBCD, 5.6 mg; A), AZ67 exposure before ACh stimulation (B), and AZ67 addition after establishment of ACh-stimulated activity (C). Glucose was maintained at 8 mM. AZ67 concentrations were 140 µg/mL (307 µM) in B and 350 µg/mL (768 µM) in C. Blue and yellow vertical ribbons identify β-cell and acinar-cell recordings, respectively. Panels include time-resolved event-halfwidth distributions, representative single-ROI traces and multi-ROI event rasters. Time and halfwidth are expressed in seconds. Color scales indicate event counts per bin. (D, E) Percentage change in β-cell event frequency (D) and event halfwidth (E) from preceding ACh-stimulated period after addition of vehicle, AZ67, or epinephrine. Underlying frequency and halwidth measurements are expressed as events min^−1^ ROI^−1^ and seconds respectively. Box plots show medians and interquartile ranges, and points represent individual islets. Dashed lines indicate no change. Pairwise comparisons used two-sided Mann–Whitney U tests with Holm correction. Adjusted p-values are shown in D; no significant between-group differences were detected in E. Each group comprised 3 mice, with 5 vehicle slices and 4 slices each for AZ67 and epinephrine.

We next asked whether preincubation with the PFKFB3 inhibitor AZ67 modifies ACh responsiveness in WT tissue. AZ67 had no detectable independent effect on β-cell activity at 8 mM glucose, and ACh-induced potentiation remained intact. Likewise, AZ67 did not measurably alter acinar-cell activity at 8 mM glucose, and ACh elicited a robust response comparable to control conditions (Fig. 4B, D). Finally, we applied AZ67 acutely after establishing ACh-driven activity in both compartments. Under these conditions, AZ67 did not produce a discernible change in β-cell oscillation frequency, whereas in acinar cells it partially attenuated the time-dependent decline in Ca^2+^ oscillatory activity (Fig. 4C).

To quantitatively benchmark the acute effect of AZ67 in WT β-cells, we compared it with vehicle and with epinephrine, a well-established physiological inhibitory input to β-cells. For each experiment, the change from the preceding ACh-stimulated period was calculated for Ca²⁺ event frequency and event halfwidth (Fig. 4D, E). Vehicle and AZ67 produced relatively small and variable changes in both parameters and did not differ significantly from one another. Epinephrine produced a large reduction in event frequency. The difference was significant relative to vehicle (Holm-adjusted p = 0.048), whereas the comparison with AZ67 did not reach the 0.05 threshold (p = 0.057). No significant between-treatment differences in event halfwidth were detected (Fig. 4D,E). Thus, the WT preparation remained readily capable of expressing a strong inhibitory Ca²⁺ phenotype, but AZ67 did not reproduce this response. These data argue against nonspecific suppression of healthy β-cell Ca²⁺ activity by AZ67 and support a context-dependent action of PFKFB3 inhibition.

### PFKFB3 inhibition enhances ACh-responsive β-cell activity and alters acinar cell oscillation dynamics in 3xTg-AD slices

We next pooled experiments in which 3xTg-AD pancreatic slices were sequentially exposed to 8 mM glucose, 100 nM ACh, and either AZ67 or the second PFKFB3 inhibitor AZ26 (Fig. 5). A representative AZ67 experiment illustrates the distinct responses of β-cells and acinar cells within the same tissue preparation (Fig. 5A).

**Figure 5.**
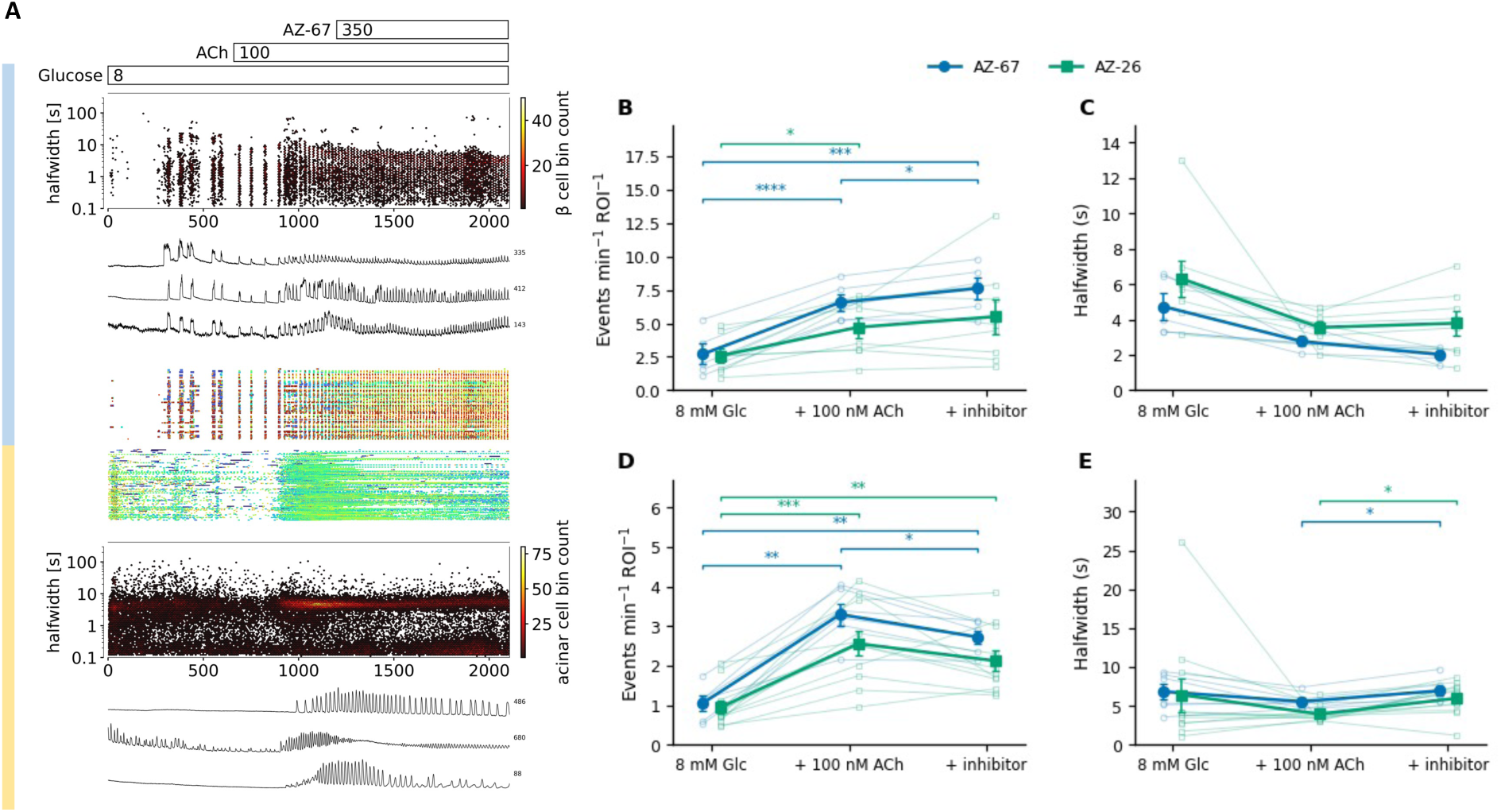
PFKFB3 inhibitors alter β-cells and acinar cell Ca^2+^ dynamics in 3xTg-AD pancreatic slices. (A) Representative recording during sequential exposure to 8 mM glucose, 100 nM ACh and AZ67. Blue and yellow vertical ribbons identify β-cell and acinar-cell recordings, respectively. Time-resolved event-halfwidth distributions, representative single-ROI traces and multi-ROI event rasters are shown. Time and halfwidth are expressed in seconds. Color scales indicate event counts per bin. (B–E) β-cell event frequency (B) and halfwidth (C), and acinar cell event frequency (D) and halfwidth (E), across the sequential conditions. Frequency is expressed as events min^−1^ ROI^−1^ and halfwidth in seconds. AZ67 (blue) or AZ26 (green) were each applied at 350 µg mL^−1^, corresponding to 768 and 870 µM, respectively, while ACh and glucose remained present. Thin lines connect individual slices. Bold symbols, lines and error bars show mean ± SEM. During inhibitor exposure, epoch-averaged acinar event frequency was lower and event halfwidth greater than during the preceding ACh period, despite renewed oscillations in the time-resolved recording. Treatment effects were analyzed separately within each inhibitor series using repeated-measures ANOVA and multiplicity-adjusted paired comparisons. The compounds were not directly compared. AZ67: 4 mice, 12 slices; AZ26: 3 mice, 11 slices. *p < 0.05, **p < 0.01, ***p < 0.001, and ****p < 0.0001.

In β-cells, ACh increased Ca²⁺ event frequency from the relatively low activity observed at 8 mM glucose and simultaneously shifted activity toward shorter-duration events (Fig. 5A-C). Subsequent addition of AZ67 produced a further significant increase in event frequency, while β-cell events remained short and became still shorter on average. AZ26 produced a qualitatively similar trajectory, with increased β-cell activity during ACh stimulation that was maintained after inhibitor addition, although the magnitude and statistical significance of individual within-protocol changes differed between the two treatment series. Because AZ67 and AZ26 experiments were analyzed independently, these data do not establish a quantitative difference in efficacy between the compounds.

During sustained exposure to 100 nM ACh, acinar Ca²⁺ responses evolved over time rather than remaining stationary. An initial high-activity phase was followed by an intermediate plateau and a progressive decline toward baseline (Fig. 5A, bottom panel). Across the full analysis epochs, mean acinar event frequency was lower and event halfwidth greater during inhibitor exposure than during the preceding ACh period (Fig. 5D,E). These epoch averages coexist with the re-emergence of oscillations visible in the time-resolved recordings. We therefore describe this response as re-establishment of oscillatory signaling rather than simply attenuation of response rundown.

### β-cell Ca^2+^ activity increases during sequential nanomolar AZ67 exposure

The high nominal AZ67 concentration used in the initial acute-rescue experiments did not establish the concentration range at which the β-cell effect first becomes detectable. We therefore exposed ACh-stimulated 3xTg-AD slices sequentially to AZ67 in the nanomolar range while following β-cell and acinar-cell Ca²⁺ dynamics in parallel (Fig. 6). β-cell Ca^2+^ event AUC increased modestly after addition of 100 nM ACh and increased further during subsequent AZ67 exposure, reaching statistical significance at 100 nM AZ67 relative to 8 mM glucose alone (Fig. 6B). Thus, the response was characterized by early recruitment followed by an apparent plateau rather than by a continuously increasing concentration-response. The increase in β-cell activity was accompanied by a marked shortening of Ca²⁺ events, after which event halfwidth remained comparatively stable across the AZ67 ramp (Fig. 6C). Together, these changes indicate recruitment of a faster oscillatory β-cell phenotype rather than prolongation of individual Ca²⁺ events.

**Figure 6.**
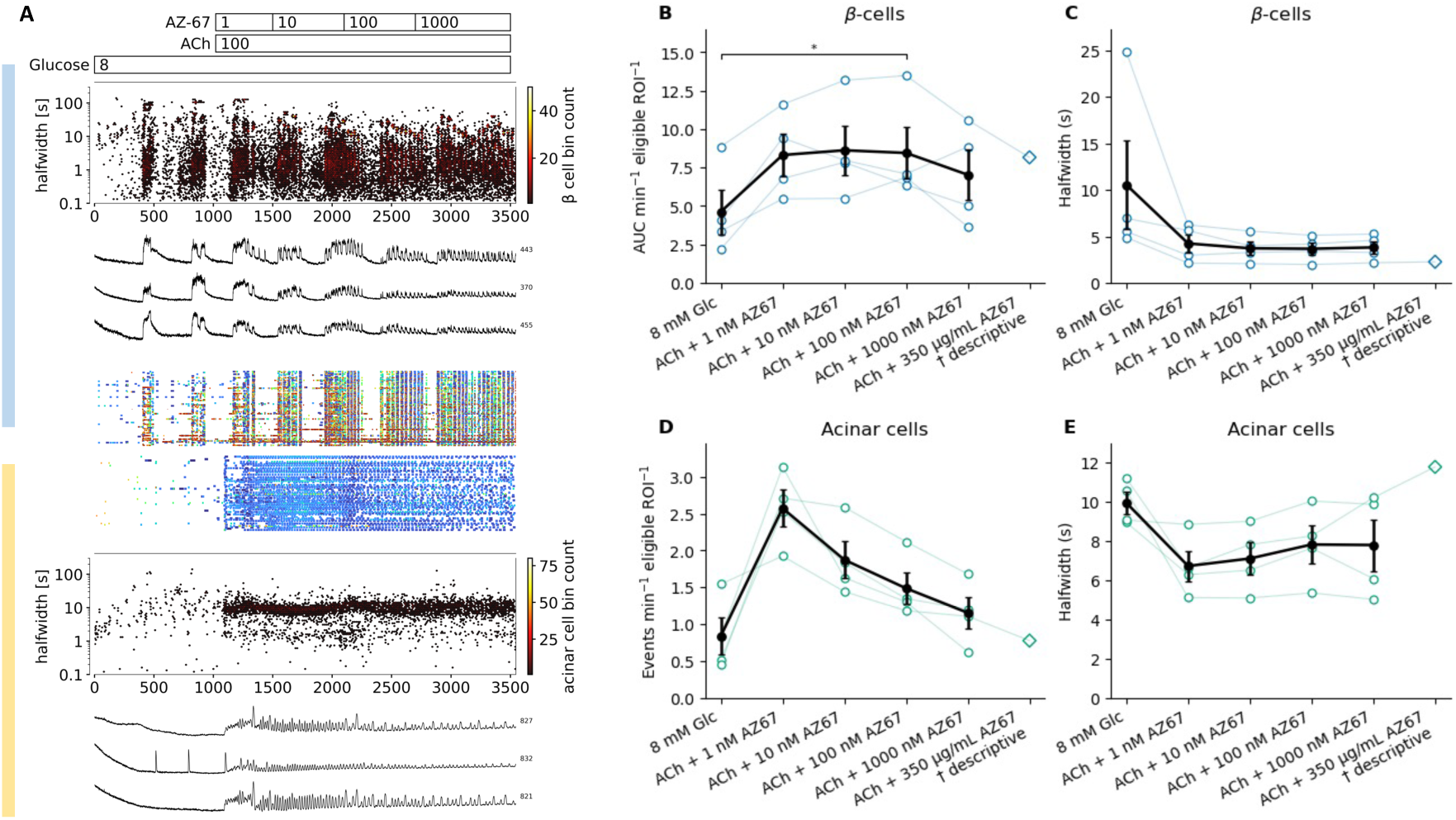
β-cell Ca^2+^ activity during sequential nanomolar AZ67 exposure. (A) Representative 3xTg-AD pancreatic tissue slice recording during stimulation with 8 mM glucose and 100 nM ACh, followed by a sequentially exposure to 1, 10, 100 and 1000 nM AZ67. Blue and yellow vertical ribbons identify β-cell and acinar-cell recordings, respectively. Time-resolved event-halfwidth distributions, representative single-ROI traces and multi-ROI event rasters are shown. Time and halfwidth are expressed in seconds. Color scales indicate event counts per bin. (B,C) β-cell event AUC per min per eligible ROI (B) and event halfwidth in seconds (C). (D, E) Acinar-cell event frequency, expressed as events min^−1^ ROI^−1^ (D) and event halfwidth in seconds (E). Open circles and thin lines show individual slices. Black symbols and error bars indicate mean ± SEM. The separate diamond denotes the descriptive high-concentration, 350 µg mL^−1^ AZ67, available in one recording and excluded from inferential statistics. Nanomolar series: 2 mice, 4 slices. Conditions shared by all recordings were compared by repeated-measures ANOVA followed by two-sided paired t-tests with Holm correction within each panel. The bracket in B indicates the comparison between 8 mM glucose alone and ACh plus 100 nM AZ67. *p < 0.05.

Acinar cells displayed a different temporal profile during the same protocol. Event frequency increased during ACh stimulation and the initial low-concentration phase but subsequently declined as the sequential exposure continued (Fig. 6D), consistent with the time-dependent rundown of ACh-evoked acinar activity observed in the other protocols. Acinar event halfwidth decreased initially and then partially recovered during later steps (Fig. 6E).

In these paired measurements the β-cell response reached statistical significance during the 100 nM AZ67 condition, while endocrine and exocrine compartments exhibited distinct responses during continued exposure.

### Adrenergic-challenge experiments suggest preserved β-cell recruitment after AZ67 exposure

We next asked whether the AZ67-restored β-cell state remained responsive in the presence of an opposing adrenergic signal. In age-matched WT β-cells, ACh increased Ca^2+^ event frequency above that observed at 8 mM glucose, whereas subsequent addition of 10 nM epinephrine strongly suppressed the ACh-potentiated response (Fig. 7C). 3xTg-AD β-cells showed lower activity than WT under both 8 mM glucose and ACh stimulation, consistent with the impaired responsiveness described above, and epinephrine further reduced their event frequency (Fig. 7A,C).

**Figure 7.**
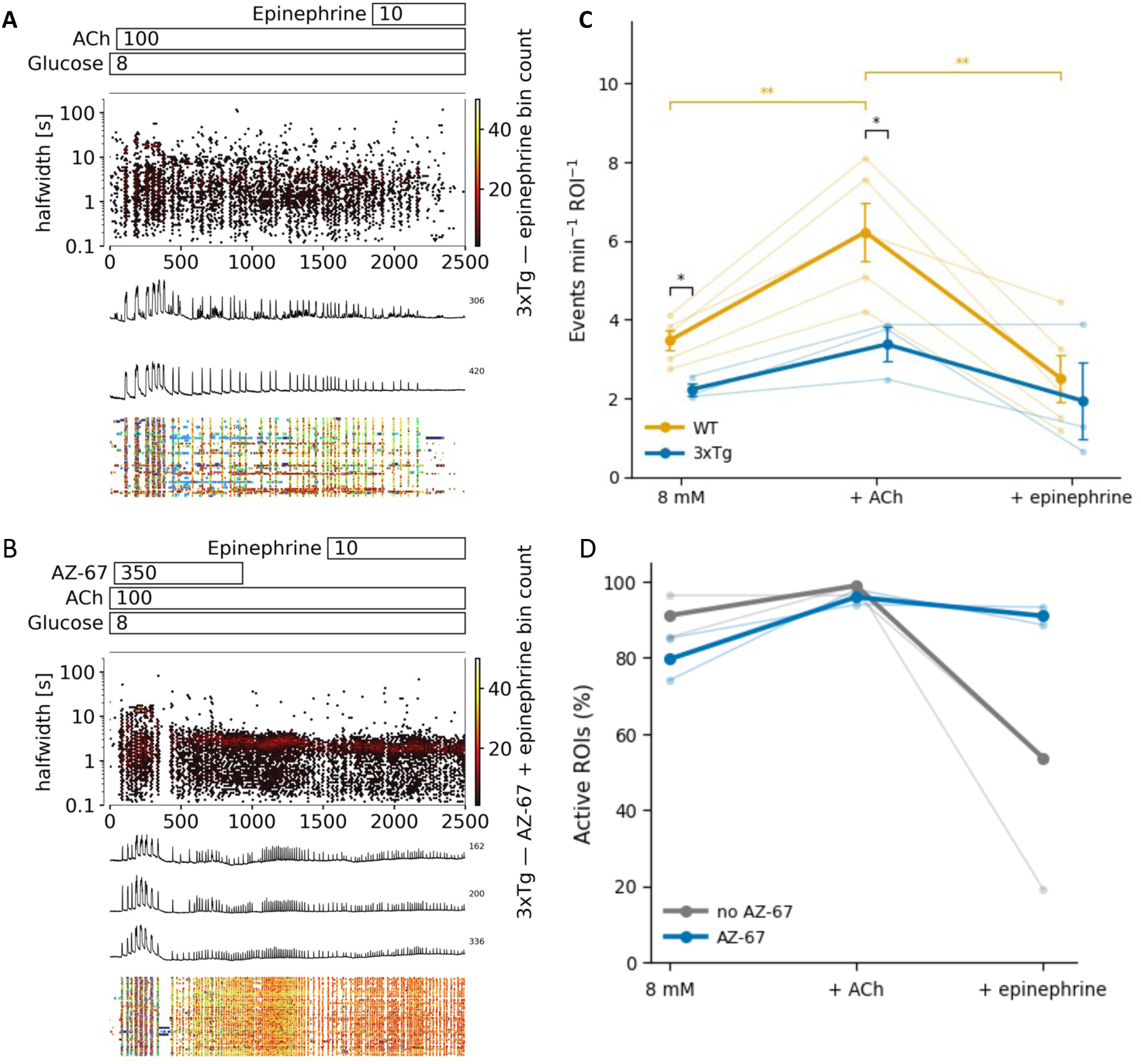
β-cell recruitment during adrenergic challenge after AZ67 exposure. (A,B) Representative 3xTg-AD pancreatic-slice recordings during stimulation with 8 mM glucose and 100 nM ACh, followed by 10 nM epinephrine, without prior AZ67 exposure (A) or after exposure to 350 µgmL^−1^ AZ67 (768 µM; B). Protocol bars indicate application intervals. Panels show time-resolved event-halfwidth distributions, representative single-ROI traces and multi-ROI event rasters. Time and halfwidth are expressed in seconds. Color scales indicate event counts per bin. (C) Ca^2+^ event frequency in active β-cell ROIs, expressed as events min^−1^ ROI^−1^, in WT (orange; 2 mice, 5 slices) and 3xTg-AD slices without AZ67 (blue; 1 mouse, 3 slices). Thin lines connect individual experiments. Bold symbols, lines and error bars indicate mean ± SEM. Orange brackets denote within-WT comparisons and black brackets denote between-genotype comparisons at the indicated conditions. (D) Active β-cell ROIs as a percentage of eligible ROIs during each protocol phase, without AZ67 exposure (grey) or following AZ67 exposure (blue). Thin lines show individual experiments and bold lines show group means; 2 mice, 2 slices per condition. *p < 0.05; **p < 0.01).

In matched 3xTg-AD experiments, the effect of epinephrine on β-cell recruitment depended markedly on PFKFB3 inhibition. Without AZ67, epinephrine caused pronounced de-recruitment of β-cells following ACh stimulation, with the fraction of active ROIs falling substantially in both experiments examined (Fig. 7A,D). In contrast, when AZ67 was present, approximately 90% of eligible β-cell ROIs remained active during epinephrine exposure (Fig. 7B,D).

In these exploratory recordings, AZ67 exposure was associated with preserved ACh-dependent Ca^2+^ activity in 3xTg-AD β-cells and with preservation of β-cell recruitment during subsequent adrenergic challenge.

### Long-duration recordings show altered β-cell activity after removal of AZ67 or AZ26 from the perfusate

The preservation of β-cell recruitment during epinephrine challenge suggested that PFKFB3 inhibition may stabilize a responsive β-cell state rather than simply producing an acute increase in Ca^2+^ activity. We therefore examined two long-duration sequential protocols in 3xTg-AD pancreatic slices to determine how β-cell and acinar-cell activity evolved during and after exposure to PFKFB3 inhibitors (Fig. 8).

**Figure 8.**
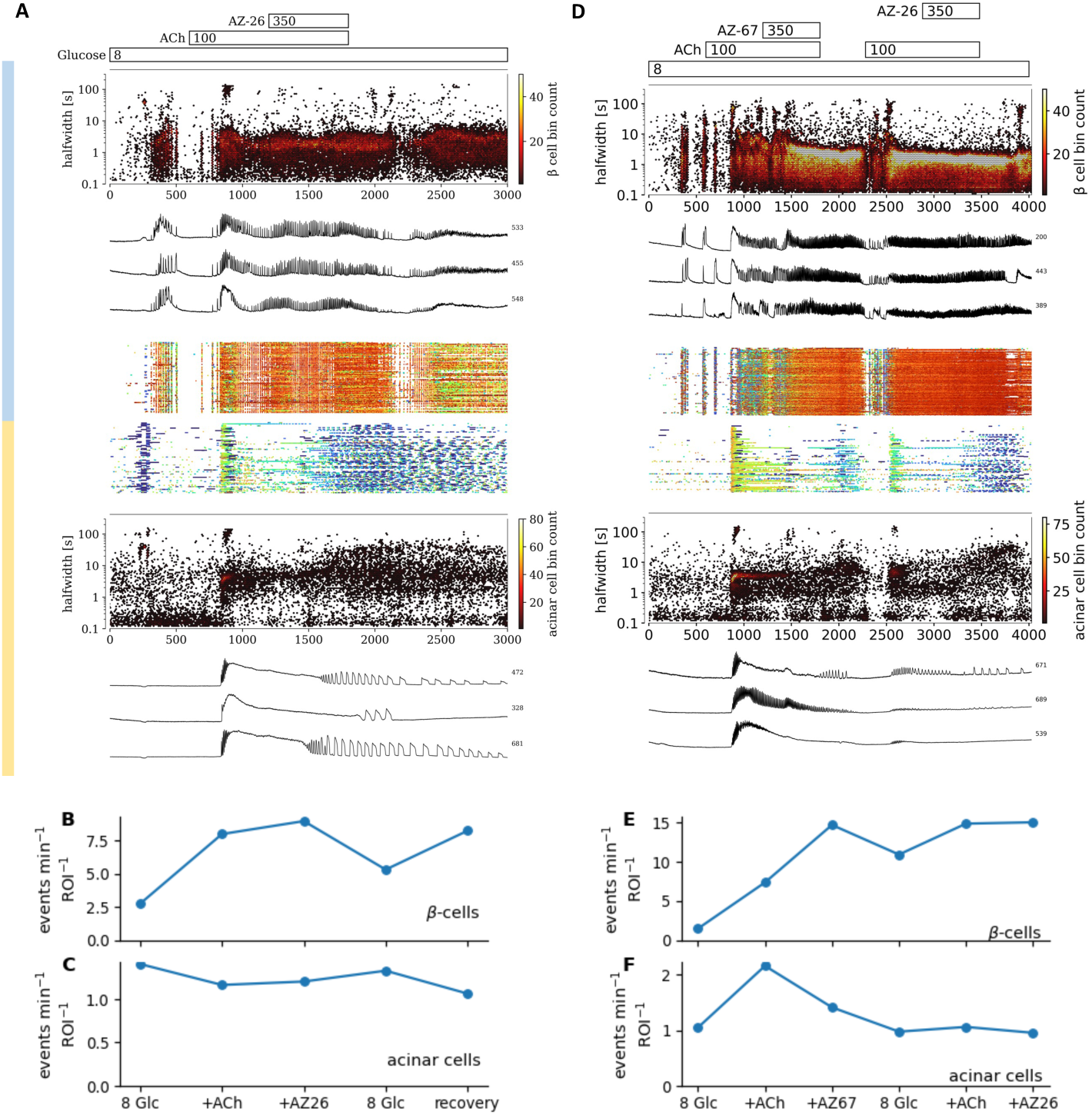
β-cell and acinar cell Ca^2+^ activity during and after PFKFB3 inhibitor exposure. (A) Representative long-duration 3xTg-AD pancreatic slice recording during sequential exposure to 8 mM glucose, 100 nM ACh and AZ26, followed by withdrawal of ACh and AZ26 while glucose was maintained. (B,C) Time-resolved Ca^2+^ event plots, representative single-cell traces and multi-ROI activity rasters are shown for β-cells (upper panels) and acinar cells (lower panels). (B,C) Corresponding event frequency in active β-cell (B) and acinar-cell (C) ROIs during glucose stimulation, ACh stimulation, AZ26 exposure, the early post-withdrawal period and later recovery. (D) Representative recording during sequential ACh and AZ67 exposure, withdrawal of both compounds, ACh re-application and subsequent AZ26 exposure at constant 8 mM glucose. (E,F) Corresponding event frequency in active β-cell (E) and acinar-cell (F) ROIs. AZ67 and AZ26 were each applied at 350 µg mL^−1^, corresponding to 768 and 870 µM, respectively. In A and D, blue and yellow vertical ribbons identify β-cell and acinar-cell recordings. Time-resolved event-halfwidth distributions, representative single-ROI traces and multi-ROI event rasters are shown; time and halfwidth are expressed in seconds, and color scales indicate event counts per bin. In B,C,E,F, frequency is expressed as events min^−1^ active ROI^−1^. Each line summarizes one representative recording; no inferential statistics were applied. Withdrawal denotes removal from the perfusate and does not establish complete clearance from the tissue.

In the first experiment, ACh increased β-cell Ca^2+^ event frequency above that observed at 8 mM glucose, and addition of AZ26 further increased the frequency of events in active β-cell ROIs (Fig. 8A,B). Following withdrawal of both ACh and AZ26 and return to 8 mM glucose alone, β-cell event frequency initially decreased but subsequently increased again during the later recovery period despite the absence of AZ26 from the perfusate. Acinar-cell event frequency changed comparatively little during the same sequence and remained low during the post-inhibitor period (Fig. 8A,C).

A second long-duration experiment tested sequential AZ67 and AZ26 exposure (Fig. 8D–F). ACh increased β-cell event frequency, and subsequent AZ67 exposure produced a further pronounced increase. Removal of ACh and AZ67 reduced β-cell activity, but event frequency remained above the initial 8 mM glucose level. Re-application of ACh in the absence of AZ67 in the perfusate rapidly restored the high-frequency β-cell phenotype, and subsequent addition of AZ26 produced no further increase (Fig. 8D,E). In contrast, ACh initially elicited robust acinar-cell activity, which progressively waned during sustained stimulation. Addition of AZ67 was followed by renewed oscillatory activity, although mean event frequency across the AZ67 exposure period remained lower than during the preceding ACh epoch. Following washout, acinar activity returned toward the initial glucose-stimulated level. During the second ACh exposure, activity again remained comparatively low until subsequent AZ26 application was accompanied by re-emergence of oscillatory activity (Fig. 8D,F).

Thus, both long-duration recordings are consistent with a β-cell functional effect that extends beyond the period of acute PFKFB3 inhibitor application. The distinct trajectories of β-cells and acinar cells further argue against a nonspecific persistent elevation of Ca^2+^ activity throughout the tissue. Because these experiments were designed as long-duration concept recordings rather than replicated washout studies, however, they do not distinguish a persistent change in β-cell functional state from prolonged tissue retention or target engagement by the inhibitors.

## DISCUSSION

Four findings emerge from these experiments. First, pharmacological PFKFB3 inhibition acutely enhances β-cell Ca²⁺ activity in 3xTg-AD pancreatic tissue slices, with onset within minutes. The functional deficit in this model seems to be modifiable, on a timescale that excludes β-cell replacement, proliferation or tissue remodeling. Second, the effect appears at nanomolar exposure, reaching significance at 100 nM AZ67, and the structurally distinct inhibitor AZ26 reproduces its direction. Third, and most consequentially, the enhanced phenotype outlasts the inhibitor and resists physiological brake, as β-cell activity remained above baseline after AZ67 and AZ26 were removed from the perifusate, ACh re-application without inhibitor restored the high-frequency phenotype, and most eligible β-cell ROIs remained active during epinephrine exposure. An effect that survives both drug removal and adrenergic input is the signature of a changed β-cell state rather than transient pharmacological stimulation and it is the property on which therapeutic value depends. Fourth, the response is compartment-specific. β-cells and neighboring acinar cells in the same slice, under identical perfusion, follow opposite trajectories.

Persistence is what gives these observations translational weight. When ACh and inhibitor were withdrawn together, β-cell event frequency fell but settled above the initial glucose-stimulated level; re-applying acetylcholine alone restored the high-frequency phenotype, and adding AZ26 gained nothing further. The tissue behaved as though it had been altered rather than merely stimulated. Acinar activity over the same sequence returned to its starting level, without a tissue-wide drift. Under adrenergic challenge with epinephrine acting through α₂-adrenergic signalling^29^, the physiological brake on insulin release, the recruitment in AZ67-treated slices was not suppressed. The therapeutic significance is direct. A compound whose benefit ends with its clearance must be given continuously and can be opposed by the counter-regulatory signaling that rises in diabetes; one that improves the β-cell does neither. This distinction may extend to durable disease modification, since the companion study by Raval et al.^18^ draws at the level of the animal, where glycemic benefit persisted after AZ67 withdrawal and where improved glucose tolerance and insulin secretion in a diabetic hIAPP model were accompanied by functional effects in stressed human islet microtissues. Our recordings locate the same property in single β-cells within intact tissue, on a timescale of minutes rather than weeks. Two qualifications bound the claim without diminishing it. Removal from the perifusate is not pharmacological washout, so residual tissue retention or prolonged target engagement remain open until replicated washout and rechallenge experiments are coupled to tissue drug measurement. And recruitment, oscillation frequency and insulin secretion are distinct outcomes: preserved recruitment under adrenergic tone is not preserved secretion.

In β-cells, inhibitor exposure recruited a faster oscillatory phenotype: event frequency rose and individual events shortened. In acinar cells recorded simultaneously, mean event frequency fell and events lengthened relative to the preceding ACh period. These are not the same response at different magnitudes; they are opposite in sign on both measures. A perfusion artefact, a pH or temperature drift, or a generalized excitability change would move both compartments together, and none does. The compartment specificity is therefore the strongest internal evidence that the β-cell response reflects cell-type specific biology rather than a nonspecific elevation of Ca²⁺ activity across the tissue.

The acinar trajectory is itself structured. Sustained ACh stimulation produces a characteristic rundown of acinar activity, and inhibitor application during that rundown was followed by renewed oscillations in the time-resolved recordings, even though epoch-averaged frequency remained below the preceding ACh period and halfwidth remained longer. The pattern applies to a reorganization of acinar Ca^2+^ dynamics with oscillatory signaling re-emerging on a lower-frequency and longer-event background. This matters because PFKFB3-dependent IP_3R_ signaling has been linked to acinar Ca^2+^overload and pancreatitis^13^. Longer events could indicate either reduced or redistributed Ca^2+^ burden, and neither frequency nor duration alone resolves the question. Thus, integrated Ca^2+^ measurements and direct injury endpoints are required before any further interpretation.

A mechanistic hypothesis follows from the timescale and from the ACh concentration-response. Glycolytic enzymes modulate IP_3 r_eceptor gating, and PFKFB3 regulates IP_3R_-dependent Ca^2+^ signaling in acinar cells^13^, making muscarinic/IP_3-_linked Ca^2+^ release a plausible point of action in stressed β-cells. The bell-shaped ACh response supports the idea that increasing muscarinic stimulation beyond the concentration giving maximal event frequency reduced oscillatory activity, so more receptor stimulation does not translate into more oscillations. If PFKFB3 upregulation contributes to excessive or poorly coordinated IP_3R_ activity, inhibition could restore oscillatory signaling by returning Ca^2+^ release toward a more favorable operating range. A genetic precedent exists in the same model background, as reducing IP_3R_1 expression normalized exaggerated neuronal Ca^2+^signalling in PS1 M146V knock-in and 3xTg mice and attenuated pathology in the latter^35^. That precedent motivates the hypothesis but does not establish that neuronal and β-cell mechanisms are identical, and direct regulation of β-cell IP_3R_s remains to be demonstrated.

The immunostaining places these responses in the context of a PFKFB3-associated stress state. Dysfunctional β-cells under diabetogenic stress upregulate PFKFB3 alongside HIF1α activation, MAIP1 expression and impaired proinsulin processing^14,15,36^, and we observed increased nuclear PFKFB3 immunoreactivity in 3xTg-AD islets with hTG+HFD tissue as positive control. Increased PFKFB3 transcript abundance has also been reported in postmortem visual cortex from people with AD^37^, although a tissue-level association does not establish a shared cellular mechanism. We therefore treat nuclear PFKFB3 as a stress-state marker rather than as the effector pool underlying the acute Ca^2+^ response, and note that differences in insulin immunoreactivity alone do not establish β-cell loss or loss of identity. Of note, although the cytoplasmic PFKFB3 pool was comparable between wild-type and 3xTg-AD mice, the diabetogenic stress arising from increasing insulin resistance may confer greater sensitivity to PFKFB3-dependent modulation of IP₃R signaling in 3xTg-AD β-cells.

The scope of these conclusions is bounded in three ways. Some experiments included few animals with multiple preparations per animal, which limits generalized statements and requires the nested design to be accounted for analysis, although our previous work supports sampling multiple preparations to capture within-animal variability^30,38^. The datasets were not designed to resolve sex-specific effects, and the ACh ramp in male WT mice provides a mechanistic framework rather than a genotype comparison. Most fundamentally, Ca^2+^ imaging does not establish improved insulin secretion in the same preparations; the sequential concentration ramp confounds concentration with exposure time and cannot define an equilibrium concentration-response relationship. The directional agreement between AZ67 and AZ26 is supporting pharmacological evidence and proof of target, in line with the target-engagement demonstrated in the companion study.

What these experiments contribute is a relevant translational entry point. They identify a component of β-cell Ca²⁺dysfunction that responds within minutes to PFKFB3 inhibition with a small molecule, at concentrations where the effect first becomes detectable, that outlasts the compound and withstands inhibitory adrenergic^18^ input, and that is confined to the endocrine compartment in tissue where endocrine and exocrine cells can be compared directly. The immediate questions are whether the persistence reflects durable reprogramming or prolonged target engagement, whether the acute response and the durable metabolic benefit reported in the companion study show multiple PFKFB3 related mechanisms, whether the acinar reorganization reduces or merely redistributes Ca^2+^ burden, and whether the IP_3R_ hypothesis survives the future low-ACh test.

## MATERIALS AND METHODS

### Animals, ethics, and pancreatic tissue slices

The majority of experiments were performed in 10–15-month-old 3xTg-AD mice of both sexes and age-matched B6129SF2 wild-type (WT) controls. ACh concentration-response experiments we performed in male C57BL/6J of similar age. 3xTg-AD and corresponding WT mice^22^ were obtained from The Jackson Laboratories (JAX), United States of America, and subsequently bred as described^21^ and aged in individually ventilated cages at 22–24 °C and 45–55% relative humidity with a 14h:10h light/dark cycle and access to food (standard breeding chow, Ssniff V1124 pellets) and water ad libitum. Prior to tissue section mice were maintained on standard chow under a 12-h light/dark cycle for up to one week. C57BL/6J mice were maintained on standard chow. Animals were housed in individually ventilated cages (Allentown LLC, USA) at 22–24 °C and 45–55% relative humidity under a 12-h light/dark cycle, with *ad libitum* access to food and water. Procedures complied with national and European regulations and were approved by the Administration of the Ministry of Education, Science and Research, Republic of Austria (BMWFW-66.007/0032-V/3b/2019, GZ 2022-0.325.009, and GZ 2023-0.691.200).

Acute pancreatic tissue slices were prepared as previously described^30,38^. Briefly, after killing by CO₂ application and cervical dislocation, the common bile duct was clamped distally and injected with 1.9% low-melting-point agarose (Lonza) in extracellular solution (ECS; in mM: 125 NaCl, 26 NaHCO₃, 6 glucose, 6 lactic acid, 3 myo-inositol, 2.5 KCl, 2 Na-pyruvate, 2 CaCl₂, 1.25 NaH₂PO₄, 1 MgCl₂, and 0.5 ascorbic acid). The pancreas was cooled in ice-cold, carbogen bubbled ECS, removed, embedded in agarose, and sectioned at 140 µm using a VT1000S vibratome (Leica Microsystems).

Slices were stored at room temperature in HEPES-buffered saline containing 6 mM glucose (in mM: 150 NaCl, 10 HEPES, 6 glucose, 5 KCl, 2 CaCl₂, and 1 MgCl₂; pH 7.4). For Ca²⁺ imaging, slices were loaded for 50 min with 6 µg Calbryte 520 AM (AAT Bioquest) in 3.33 mL HEPES-buffered saline supplemented with 3.75 µL dimethylsulfoxide and 1.25 µL Pluronic F-127.

### Ca^2+^ imaging

Slices were continuously perfused with ECS at 37 °C in a 0.3-mL recording chamber at 0.72 mL min^−1^. Imaging was performed using a Leica SP8 Falcon confocal microscope with a 20× HC PL APO CS2 objective (NA 0.75) or a Nikon Ti inverted confocal with a 20× APO LWD WI λS objective (NA 0.95). Calbryte 520 was excited at 488 nm and fluorescence collected at 500–700 nm as previously described^30^. The optical section thickness was approximately 5 µm. Images were acquired at 256 × 256 pixels and 20 Hz on the Leica or 30 Hz on the Nikon system. Pixel size was approximately 1 μm^2^.

### Stimulation Protocols

During imaging, slices were continuously perfused at 37 °C with ECS. Glucose dependence was assessed using stepwise increases from 4 to 8 mM glucose. Muscarinic potentiation was examined by adding 100 nM ACh at 8 mM glucose. For the ACh concentration-response experiments, WT slices were sequentially exposed to 0, 100, 250 and 500 nM ACh at constant 8 mM glucose. Application intervals are shown in the corresponding figure panels.

For acute inhibitor experiments, AZ67 (350 µg mL^−1^; 768 µM) or AZ26 (350 µg mL^−1^; 870 µM) was added after establishment of the ACh response, while 8 mM glucose and ACh remained present.

For the nanomolar AZ67 series, 3xTg-AD slices were exposed sequentially to 1, 10, 100, and 1000 nM AZ67 during stimulation with 8 mM glucose and 100 nM ACh. β-cell and acinar-cell activity was recorded simultaneously. A subsdquent high-concentration condition, 350 µg mL^−1^ AZ67 was available in one recording and was analyzed descriptively.

For adrenergic-challenge experiments, 10 nM epinephrine was applied during stimulation with 8 mM glucose and 100 nM ACh. Experiments were performed in WT and 3xTg-AD slices without inhibitor and in 3xTg-AD slices previously exposed to AZ67. In the latter protocol, AZ67 was removed from the perfusate before epinephrine application, as indicated by the protocol bars.

Long-duration recordings examined activity during and after inhibitor exposure at constant 8 mM glucose. In the AZ26 protocol, ACh and AZ26 were withdrawn together. In the sequential AZ67/AZ26 protocol, ACh and AZ67 were withdrawn together, followed by ACh re-application and subsequent AZ26 exposure. Withdrawal denotes removal from the perifusate, tissue drug concentrations were not measured. The longest washout interval was approximately 20 min.

### Ca^2+^ event analysis

Ca^2+^ imaging data were analyzed using previously described Python-based workflows^30^. Cellular regions of interest (ROIs) and Ca^2+^ events were detected automatically. β-cells ROIs were identified by intra-islet localization and their characteristic glucose-dependent activity^29,30,38^. Other endocrine cell types were not included. β-cell and acinar-cell ROIs were analyzed separately.

Event halfwidth, amplitude and area under the curve (AUC) were quantified for each detected event^38^. Events with halfwidths outside 0.1–100 s were excluded. Unless otherwise specified, ROIs with fewer than five qualifying events over the complete recording were excluded. ROIs meeting this recording-wide criterion were designated eligible ROIs, and the same eligible population was used across protocol phases within each recording.

Event frequency was calculated from the number of qualifying events divided by the duration of the analysis epoch in minutes. For measurements expressed per eligible ROI, eligible ROIs without events in a given epoch contributed zero. For measurements expressed per active ROI, only ROIs with more than 5 events within that epoch were included. β-cell recruitment was calculated as the percentage of eligible ROIs meeting this epoch-specific activity criterion.

Measurements were aggregated at the recording/slice level before group comparisons; individual events and ROIs were not treated as independent biological replicates. Within each recording and analysis epoch, event halfwidth, amplitude and AUC were summarized as medians within each ROI, followed by the median across ROIs to obtain an islet-level value, while frequency was calculated from event counts, and cumulative AUC from summed event areas, before aggregation across ROIs. Long-duration recordings in Fig. 8 were analyzed descriptively over the indicated protocol epochs.

### Chemicals and Reagents

AZ67 (Cat. No. 5742) and its analogue AZ26 (Cat. No. 5675) were purchased from Tocris Biosciences. 2-hydroxypropyl-β-cyclodextrin (Cat. No. H107) were purchased from Sigma Aldrich, Millipore Sigma, USA.

### Histology and immunofluorescence

Pancreatic tissue was fixed in 4% paraformaldehyde overnight at 4°C, embedded in paraffin and sectioned at 4 µm^17^. Immunofluorescence analysis was performed using ZEN imaging software on a Zeiss Axio Imager 2.0 microscope equipped with an Apotome structured illumination module. The following primary antibodies were used with overnight incubation at 4°C : rabbit anti-PFKFB3 (Abcam ab181861, Cambridge, MA, USA, 1:100), and Insulin Monoclonal Antibody (ICBTACLS), eBioscience™ (Invitrogen, 14-9769-82; 1:200), Secondary antibodies were applied for 1h at room temperature and included Goat anti-Rabbit IgG (H+L) Highly Cross-Adsorbed Secondary Antibody, Alexa Fluor™ 647 (Thermo Scientific, A-21245; 1:100), and Goat anti-Mouse IgG (H+L) Highly Cross-Adsorbed Secondary Antibody, Alexa Fluor™ Plus 555 (Thermo Scientific, A32727; 1:100). Nuclei were counterstained and slides were mounted using VECTASHIELD antifade mounting medium containing DAPI (Vector Laboratories, H-1200). As a positive control for PFKFB3⁺ β-cell dysfunction, pancreata from h-IAPP-transgenic mice maintained on a high-fat diet (h-Tg+HFD) were processed and stained in parallel using the identical protocol described above.

Fluorescence images were acquired using ZEN (Zeiss) under identical acquisition settings and analyzed in QuPath. Islets were manually annotated, individual cells were detected, and nuclear and cytoplasmic mean fluorescence intensities for insulin and PFKFB3 were quantified. Cells were classified using automatically suggested fluorescence intensity thresholds from the tool single-cell intensity distribution, wherein the thresholds were decided based on mean, mean+1 SD and mean+ 2 SD. At the suggested thresholds, the compliance with visual cell positivity was matched with respect to high marker expressing cells and if needed thresholds were tweaked before applying to categorizing cells with different expression levels. So, based on these final thresholds, cells were categorized as INS^high^, INS^low^, or INS^−^, while nuclear and cytoplasmic PFKFB3 expression was classified as high, medium or low. The resulting data were analyzed and visualized using GraphPad Prism.

### Statistical Analysis

The independent experimental unit was the pancreatic slice/recording unless stated otherwise. Statistical tests were selected according to experimental design and distributional assumptions. Repeated measurements within the same preparation were analyzed using repeated-measures procedures with multiplicity-adjusted post hoc comparisons; independent groups were analyzed using the corresponding unpaired tests. Statistical significance was defined as *p* < 0.05.

For ACh concentration-response experiments (Fig. 2), the repeated 0, 100, 250, and 500 nM ACh conditions were analyzed by repeated-measures ANOVA, with paired two-tailed *t*-tests and Holm correction where indicated. Bell-shaped concentration dependence of event frequency was tested using a planned one-sided inverted-U contrast with the predicted maximum within the tested concentration range (100 nM).

For the acute AZ67 and AZ26 sequential experiments, treatment conditions were compared within each inhibitor series using repeated-measures ANOVA followed by multiplicity-adjusted paired post hoc tests. AZ67 and AZ26 series were not directly compared.

For Fig. 4D,E, percentage changes from the ACh-stimulated pretreatment period were calculated and treatment groups were compared using two-sided Mann–Whitney U tests with Holm correction.

Glucose dependence of Ca^2+^ AUC was analyzed using a linear mixed-effects model with genotype, glucose concentration, and their interaction as fixed effects and experiment identity as a random intercept (Python, statsmodels, v0.13.5). Models were fitted using restricted maximum likelihood and fixed effects evaluated using Wald statistics.

Histological group comparisons were performed using the tests specified in the corresponding figure legends. Data are presented as mean ± SEM unless otherwise indicated.

## DATA AVAILABILITY

All raw data, analytical pipelines, and processed datasets are available from Medical University of Vienna upon reasonable request.

## ACKNOWLEDGEMENTS

MSR was funded by Vienna Science and Technology Fund WWTF (LS23-026), KR, GB, and ST were funded by Hirshberg funding HF-2023-024 and Sponsored research agreement SRA 2021-026 with Metanoia Bio. T.E. is grateful to the Austrian Science Fund (FWF) for excellence cluster 10.55776/COE14, which was additionally supported by Land Styria, Stadt Graz and the University of Graz and the Medical Universities of Vienna and Graz. We are grateful for the assistance provided by all members of the animal facility team at IMB, Graz.

## AUTHOR CONTRIBUTIONS

M.S.R. and S.T. designed the study; M.S.R., S.P., and co-authors performed calcium imaging experiments; KR and GB performed and quantified PFKFB3 immunostaining; S.T. and M.S.R. wrote the manuscript with input from all authors.

## REFERENCES

1 Foskett, J. K., White, C., Cheung, K. H. & Mak, D. O. Inositol trisphosphate receptor Ca2+ release channels. Physiol Rev 87, 593–658 (2007). 10.1152/physrev.00035.2006

2 Streb, H., Irvine, R. F., Berridge, M. J. & Schulz, I. Release of Ca2+ from a nonmitochondrial intracellular store in pancreatic acinar cells by inositol-1,4,5-trisphosphate. Nature 306, 67–69 (1983). 10.1038/306067a0

3 Petersen, O. H. & Tepikin, A. V. Polarized calcium signaling in exocrine gland cells. Annu Rev Physiol 70, 273–299 (2008). 10.1146/annurev.physiol.70.113006.100618

4 Thorn, P., Lawrie, A. M., Smith, P. M., Gallacher, D. V. & Petersen, O. H. Local and global cytosolic Ca2+ oscillations in exocrine cells evoked by agonists and inositol trisphosphate. Cell 74, 661–668 (1993). 10.1016/0092-8674(93)90513-p

5 Blondel, O. et al. Localization of inositol trisphosphate receptor subtype 3 to insulin and somatostatin secretory granules and regulation of expression in islets and insulinoma cells. Proceedings of the National Academy of Sciences 91, 7777–7781 (1994). doi:10.1073/pnas.91.16.7777

6 Gilon, P., Nenquin, M. & Henquin, J. C. Muscarinic stimulation exerts both stimulatory and inhibitory effects on the concentration of cytoplasmic Ca2+ in the electrically excitable pancreatic B-cell. Biochem J 311 **( Pt** **1**), 259–267 (1995). 10.1042/bj3110259

7 Gautam, D. et al. A critical role for beta cell M3 muscarinic acetylcholine receptors in regulating insulin release and blood glucose homeostasis in vivo. Cell metabolism 3, 449–461 (2006). 10.1016/j.cmet.2006.04.009

8 Futatsugi, A. et al. IP3 receptor types 2 and 3 mediate exocrine secretion underlying energy metabolism. Science 309, 2232–2234 (2005). 10.1126/science.1114110

9 Seino, S., Shibasaki, T. & Minami, K. Pancreatic beta-cell signaling: toward better understanding of diabetes and its treatment. Proc Jpn Acad Ser B Phys Biol Sci 86, 563–577 (2010). 10.2183/pjab.86.563

10 Pfabe, J. et al. Calorie restriction modulates beta cell IP(3)R activity to regulate Ca(2+) homeostasis and cell network connectivity. Cell Calcium 136, 103161 (2026). 10.1016/j.ceca.2026.103161

11 Patterson, R. L., van Rossum, D. B., Kaplin, A. I., Barrow, R. K. & Snyder, S. H. Inositol 1,4,5-trisphosphate receptor/GAPDH complex augments Ca2+ release via locally derived NADH. Proc Natl Acad Sci U S A 102, 1357–1359 (2005). 10.1073/pnas.0409657102

12 Lavik, A. R. et al. A non-canonical role for pyruvate kinase M2 as a functional modulator of Ca(2+) signalling through IP(3) receptors. Biochim Biophys Acta Mol Cell Res 1869, 119206 (2022). 10.1016/j.bbamcr.2021.119206

13 Zhang, T. et al. PFKFB3 controls acinar IP3R-mediated Ca2+ overload to regulate acute pancreatitis severity. JCI Insight 9 (2024). 10.1172/jci.insight.169481

14 Montemurro, C. et al. IAPP toxicity activates HIF1alpha/PFKFB3 signaling delaying beta-cell loss at the expense of beta-cell function. Nat Commun 10, 2679 (2019). 10.1038/s41467-019-10444-1

15 Nomoto, H. et al. Activation of the HIF1alpha/PFKFB3 stress response pathway in beta cells in type 1 diabetes. Diabetologia 63, 149–161 (2020). 10.1007/s00125-019-05030-5

16 Min, J. et al. β-cell-specific deletion of PFKFB3 restores cell fitness competition and physiological replication under diabetogenic stress. Communications Biology 5, 248 (2022). 10.1038/s42003-022-03209-y

17 Raval, K. et al. Dysfunctional beta-cell longevity in diabetes relies on energy conservation and positive epistasis. Life Sci Alliance 7 (2024). 10.26508/lsa.202402743

18 Raval, K. et al. Targeting PFKFB3 Metabolic Gatekeeping of β-Cell Dysfunctional States Enables Durable Glucose Tolerance in Type 2 Diabetes. bioRxiv, 2026.2009.2020.752954 (2026). 10.64898/2026.09.20.752954

19 Boyd, S. et al. Structure-Based Design of Potent and Selective Inhibitors of the Metabolic Kinase PFKFB3. J Med Chem 58, 3611–3625 (2015). 10.1021/acs.jmedchem.5b00352

20 Burmistrova, O. et al. Targeting PFKFB3 alleviates cerebral ischemia-reperfusion injury in mice. Scientific reports 9, 11670 (2019). 10.1038/s41598-019-48196-z

21 Koshenov, Z. et al. Presenilin-1 controls glycolysis and identity of pancreatic beta cells. Commun Biol (2026). 10.1038/s42003-026-10581-6

22 Oddo, S. et al. Triple-transgenic model of Alzheimer’s disease with plaques and tangles: intracellular Abeta and synaptic dysfunction. Neuron 39, 409–421 (2003). 10.1016/s0896-6273(03)00434-3

23 Hamilton, L. K. et al. Central inhibition of stearoyl-CoA desaturase has minimal effects on the peripheral metabolic symptoms of the 3xTg Alzheimer’s disease mouse model. Scientific reports 14, 7742 (2024). 10.1038/s41598-024-58272-8

24 Robison, L. S. et al. Role of sex and high-fat diet in metabolic and hypothalamic disturbances in the 3xTg-AD mouse model of Alzheimer’s disease. J Neuroinflammation 17, 285 (2020). 10.1186/s12974-020-01956-5

25 Vandal, M. et al. Age-dependent impairment of glucose tolerance in the 3xTg-AD mouse model of Alzheimer’s disease. Faseb j 29, 4273–4284 (2015). 10.1096/fj.14-268482

26 Caberlotto, L. et al. Cross-disease analysis of Alzheimer’s disease and type-2 Diabetes highlights the role of autophagy in the pathophysiology of two highly comorbid diseases. Scientific reports 9, 3965 (2019). 10.1038/s41598-019-39828-5

27 Krotee, P. et al. Common fibrillar spines of amyloid-beta and human islet amyloid polypeptide revealed by microelectron diffraction and structure-based inhibitors. J Biol Chem 293, 2888–2902 (2018). 10.1074/jbc.M117.806109

28 Oskarsson, M. E. et al. In vivo seeding and cross-seeding of localized amyloidosis: a molecular link between type 2 diabetes and Alzheimer disease. Am J Pathol 185, 834–846 (2015). 10.1016/j.ajpath.2014.11.016

29 Sluga, N. et al. Physiological levels of adrenaline fail to stop pancreatic beta cell activity at unphysiologically high glucose levels. Front Endocrinol (Lausanne) 13, 1013697 (2022). 10.3389/fendo.2022.1013697

30 Postić, S. et al. Tracking Ca2+ dynamics in NOD mouse islets during spontaneous diabetes development. Diabetes (2023). 10.2337/db22-0952

31 Stožer, A. et al. Confocal Laser Scanning Microscopy of Calcium Dynamics in Acute Mouse Pancreatic Tissue Slices. J Vis Exp (2021). 10.3791/62293

32 Sluga, N. et al. Dual Mode of Action of Acetylcholine on Cytosolic Calcium Oscillations in Pancreatic Beta and Acinar Cells In Situ. Cells 10 (2021). 10.3390/cells10071580

33 Talchai, C., Xuan, S., Lin, H. V., Sussel, L. & Accili, D. Pancreatic beta cell dedifferentiation as a mechanism of diabetic beta cell failure. Cell 150, 1223–1234 (2012). 10.1016/j.cell.2012.07.029

34 Puri, S., Akiyama, H. & Hebrok, M. VHL-mediated disruption of Sox9 activity compromises β-cell identity and results in diabetes mellitus. Genes Dev 27, 2563–2575 (2013). 10.1101/gad.227785.113

35 Shilling, D. et al. Suppression of InsP3 receptor-mediated Ca2+ signaling alleviates mutant presenilin-linked familial Alzheimer’s disease pathogenesis. J Neurosci 34, 6910–6923 (2014). 10.1523/JNEUROSCI.5441-13.2014

36 Raval, K. et al. Dysfunctional β-cell longevity in diabetes relies on energy conservation and positive epistasis. bioRxiv, 2024.2003.2018.585508 (2024). 10.1101/2024.03.18.585508

37 Mjaaseth, U. N., Hsu, M. F., Arballo, J., Haj, F. G. & Patel, V. Dysregulated glucose metabolism in the visual cortex of human subjects with mild cognitive impairment and Alzheimer’s disease. Front Aging Neurosci 18, 1710075 (2026). 10.3389/fnagi.2026.1710075

38 Postić, S. et al. High-resolution analysis of the cytosolic Ca(2+) events in β cell collectives in situ. Am J Physiol Endocrinol Metab 324, E42–e55 (2023). 10.1152/ajpendo.00165.2022

